# A single-chain antibody-based AID2 system for conditional degradation of GFP-tagged and untagged proteins

**DOI:** 10.1101/2025.01.21.633519

**Authors:** Moutushi Islam, Takefumi Negishi, Naomi Kitamoto, Yuki Hatoyama, Kanae Gamo, Ken-ichiro Hayashi, Masato T. Kanemaki

## Abstract

Protein knockdown using an improved auxin-inducible degron (AID2) technology has proven to be a powerful tool for studying protein function. The current approach requires the fusion of target proteins with a degron tag, a process typically achieved through CRISPR knock-in. However, knock-in remains challenging in non-model organisms and humans, limiting the broader applicability of AID2. To overcome this limitation, we developed a single-chain antibody AID2 (scAb-AID2) system. This approach employs an adaptor composed of a single-chain antibody fused with a degron, which recognises a target protein and induces rapid degradation in the presence of the inducer 5-Ph-IAA. We demonstrated that scAb-AID2, in combination with an anti-GFP nanobody, degraded GFP-fused proteins in human cells and *C. elegans*. Furthermore, we showed that endogenous p53 and H/K-RAS were conditionally degraded in cells expressing an adaptor encoding an anti-p53 nanobody and -RAS monobody, respectively, and led to aphidicolin sensitivity in cell culture and growth inhibition in mouse xenografts. This study paves the way for broader application of AID2-based target depletion in model and non-model organisms and for advancing therapeutic strategies.

## Introduction

Conditional protein depletion is a powerful strategy for investigating protein functions in biological systems. For this purpose, siRNA and other tools suppressing mRNA production have been used for a long time^1^. In all cases, target proteins may stay for a relatively long period of time, potentially complicating the resultant phenotype by the accumulation of secondary defects. To avoid this problem, it is important to deplete target proteins in a short period of time^2,3^. By leveraging the ubiquitin-proteasome pathway equipped in eukaryotic cells, protein knockdown systems enable the direct degradation of target proteins, and this approach facilitates the observation of early-stage defects. Examples of such systems include proteolysis-targeting chimaeras (PROTACs), Trim-Away and chemically induced degron systems^4–11^.

We led the establishment of the auxin-inducible degron (AID) system that allows target proteins fused with a degron tag to be degraded in an auxin-dependent manner^12^. The AID system employs a plant-specific degradation pathway controlled by the auxin receptor TIR1 (Transport Inhibitor Response 1) and the phytohormone auxin (indole-3-acetic acid; IAA)^13^. TIR1 expressed in non-plant cells forms a functional SCF–TIR1 E3 ligase complex with the endogenous SCF subunits. In this background, a target protein fused with a degron tag derived from IAA17 is recognized by the SCF–TIR1 E3 ligase in the presence of auxin^12^.

The original AID system went through several improvements. We and others identified shorter degrons: mini-AID (mAID), AID* and mini-IAA7^14–16^. TIR1 derived from *Oryza sativa* (OsTIR1) showed better performance, even at 37℃ than one from *Arabidopsis thaliana* (AtTIR1)^12^. Recently, we and others established a new version, namely AID2 (also known as ssAID), following a bump-and-hole strategy used in chemical biology^17,18^. We utilised a combination of an OsTIR1(F74G) mutant and 5-Ph-IAA as a new inducer, enabling us to suppress leaky degradation observed with the original AID system and to induce sharper target protein degradation with a 670 times lower concentration of the inducing ligand^17^. Recently, we demonstrated degrading two proteins independently or enhancing degradation by combining AID2 and BromoTag^19^. Finally, we and others have successfully applied AID2 to degrade endogenous proteins in living *C. elegans* and mice, demonstrating that AID2 is a powerful tool for target protein degradation not only in cell culture but in living organisms^20–24^. Although AID2 is a promising tool for achieving conditional degradation in cells and animals, the target proteins must be directly fused with a degron tag. To accomplish this, CRISPR–Cas9-medicated knock-in has been employed^25,26^. However, CRISPR knock-in in non-model organisms and humans remains challenging, limiting the broader applicability of AID2.

Single-chain antibodies, such as nanobodies and monobodies derived from Camelid antibody and the fibronectin type III domain, respectively, are a valuable tool in biology and therapeutics^27–30^. Previous reports have shown that nanobodies and monobodies recognise target proteins for degradation. GFP-fused proteins were degraded when a fusion E3 ligase with an anti-GFP nanobody was expressed in mammalian cells, *Drosophila* and zebrafish^31–34^. Using a similar strategy, a SHP2 nanobody and RAS monobody were utilised to degrade endogenous SHP2 and RAS, respectively^33,35^. In all cases, the E3 ligase fusions with monobody or nanobody were expressed continuously. Therefore, the temporal control and reversibility of this strategy were rather poor. For conferring a better temporal control with a chemical ligand, the antibody-based degradation systems were combined with AID or PROTAC to degrade YFP- or GFP-fused proteins, respectively^36,37^. However, these studies were mainly focusing on degrading fluorescent proteins in cell culture.

To overcome the current limitations of AID2 in the necessity of precisely fusing the degron tag to the target proteins by genome editing, we developed the single-chain antibody AID2 (scAb-AID2). Initially, we combined an anti-GFP nanobody and AID2, successfully degrading GFP-fused proteins in human cells and *C. elegans*. We demonstrated that the expression levels of the GFP-fused target and the nanobody-degron fusion are critical for achieving efficient degradation. Moreover, we extended the application of scAb-AID2 to degrade untagged endogenous proteins, such as p53 and H/K-RAS, revealing the depleted cells show phenotypic defects in cell culture and mouse xenografts.

## Results

### The principle of scAb-AID2

To overcome the limitation of tagging endogenous proteins of interest with a degron (mini-AID or mAID) using genome editing, we utilized a single-chain antibody (scAb) for target recognition as previously shown by other groups (**Fig. 1A**)^36–38^. For this purpose, we constructed an “adaptor” composed of a scAb and mAID, and expressed it together with an E3 ligase component OsTIR1(F74G) from an all-in-one vector (**Fig. 1B**). As a proof-of-principle study, we employed a well-established anti-GFP nanobody (vhhGFP4) to degrade GFP-fused proteins and constructed three adaptors; the WT, KR and Hybrid adaptors (**Fig. 1C**)^31,36^. The WT adaptor was composed of the anti-GFP nanobody and mAID, which contain 3 and 8 lysine residues, respectively (**Fig. 1C, WT adaptor**). When the degradation inducer, 5-Ph-IAA, is applied, these lysine residues within the WT adaptor are expected to be targeted for ubiquitylation, causing its premature degradation. It is essential that an ideal adaptor is stable and recognizes its target protein without being degraded. For this purpose, all lysine residues were substituted with arginine residues to avoid ubiquitylation and degradation (**Fig. 1C, KR adaptor**)^36^. The Hybrid adaptor contained the arginine substitution in mAID, but retained three lysine residues within the anti-GFP nanobody (**Fig 1C, Hybrid adaptor**). We confirmed that the WT adaptor was degraded upon the addition of the degradation inducer 5-Ph-IAA, but the KR and Hybrid adaptors were resistant (**Fig. 1D**). These results suggest that the lysine residues within the mAID are targeted for ubiquitylation, while those within the anti-GFP nanobody are not.

**Figure 1.**
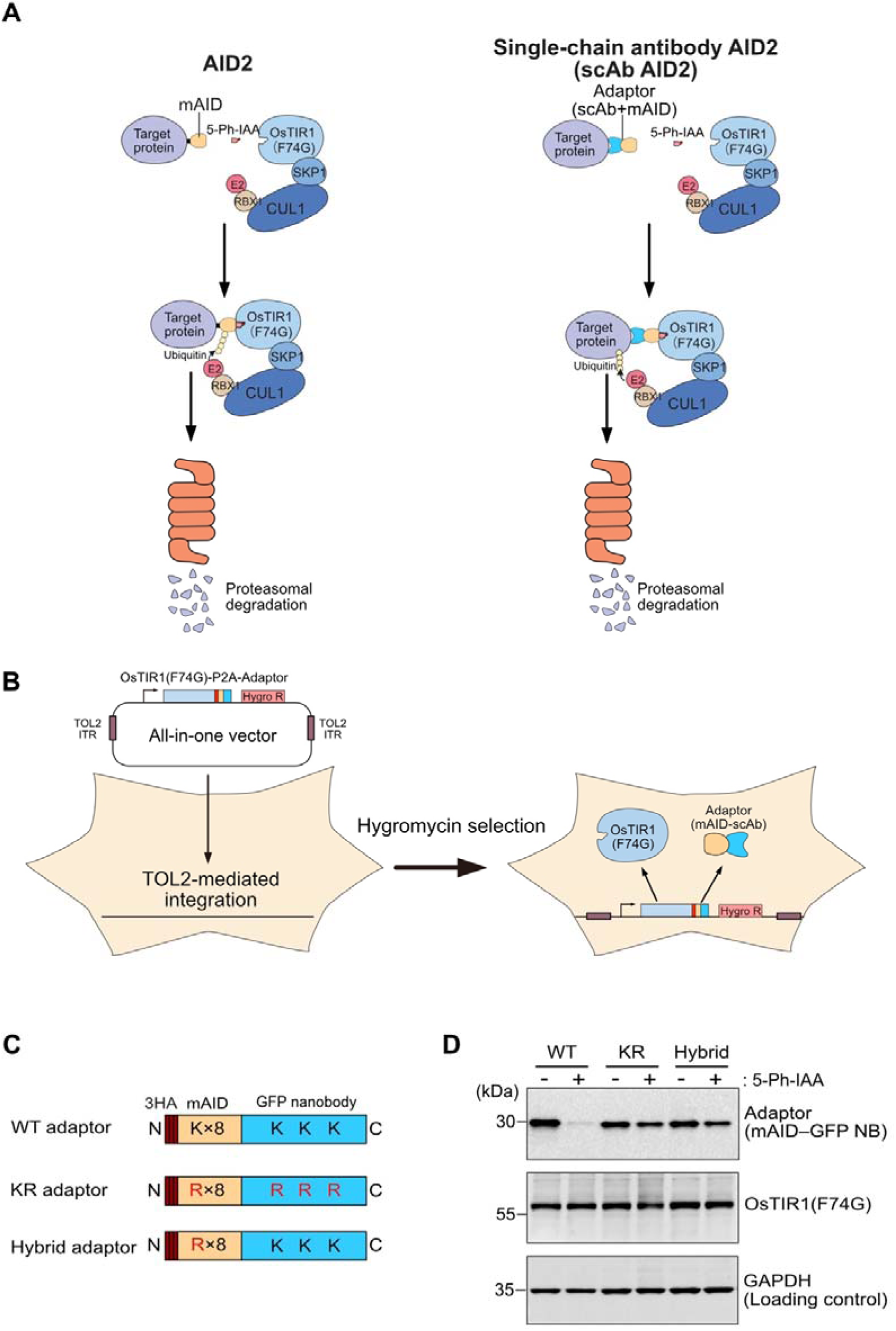
Outlines of the sc-AID2 system to degrade proteins by employing a single-chain antibody. **(A)** Schematic illustration of the AID2 and sc-AID2 systems. **(B)** Schematic illustration of an all-in-one vector for expressing OsTIR1(F74G) and adaptor. A stable cell population was selected without clone isolation. **(C)** Three types of adaptors with an anti-GFP nanobody. WT adaptor contains an unmodified degron (mAID) and the vhhGFP4 nanobody. KR adaptor contains lysine-less mAID and vhhGFP4 nanobody. Hybrid adaptor contains lysine less mAID and unmodified vhhGFP4 nanobody. **(D)** Stability of the GFP nanobody adaptors. Cells transfected with an all-in-one vector were treated for 6 h with or without 5-Ph-IAA. The adaptors and OsTIR1(F74G) were detected by Western blotting using anti-HA and anti-OsTIR1 antibodies, respectively. GAPDH was used as a loading control.

### Protein knockdown of GFP-fused proteins by scAb-AID2

Next, we used an HCT116 cell line expressing SMC6 or RAD21 fused with GFP from the endogenous locus. Both SMC6 and RAD21 form a ring-like protein complex called SMC5/6 and cohesin, respectively, and play essential roles in forming chromosome architecture through their topological DNA binding^39,40^. We introduced the all-in-one vector encoding an anti-GFP nanobody adaptor and OsTIR1(F74G) and then selected stable cells without clone isolation (**Fig. 1B**). We investigated the expression levels of SMC6– and RAD21–GFP by Western blotting and a flowcytometry after the incubation with 1 µM 5-Ph-IAA for 4 h.

When we treated the cells expressing SMC6–GFP, we observed an efficient depletion of SMC6–GFP in cells expressing the KR or Hybrid adaptor (**Fig. 2A, B, KR** and **Hybrid**). In contrast, 45.3% of SMC6–GFP remained in the cells expressing the WT adaptor (**Fig. 2B, WT**), consistent with the notion that the WT adaptor was degraded (**Fig. 1D**). We next investigated the mitotic chromosomes in the cells expressing the KR adaptor and revealed that many cells exhibited lagging chromosomes and chromosome bridges (**Fig. 2C, D**), consistent with our previously reported on SMC6-depleted cells by the AID system^41^. These results indicate that SMC6–GFP was efficiently depleted by the scAb-AID2 system. In the case of RAD21–GFP, it was depleted to 40% with the KR and Hybrid adaptors (**Fig. 2E, F**). Even though RAD21–GFP was not fully depleted, we successfully observed that mitotic chromosomes lost the sister-chromatid cohesion, indicating that cohesin function was suppressed in the RAD21–GFP depleted cells as we previously reported (**Fig. 2G**)^17^. Subsequently, we tested the reversibility of scAb-AID2. For this purpose, we initially induced SMC6– or RAD21–GFP depletion by adding 50 nM 5-Ph-IAA for 4 h. Then, we washed the cells and cultured them in a fresh media without 5-Ph-IAA for 24 h (**Supplementary Fig. 1A, B**). We found that, in both cases, the expression levels of SMC6– and RAD21–GFP were fully recovered. Additionally, we tested a cytoplasmic protein and chose a component of the cytoplasmic dynein, DHC1. DHC1–GFP was depleted to 60% with the KR and Hybrid adaptors (**Supplementary Fig. 1C, D**). Again, although the depletion of DHC1–GFP was not complete, we observed a significant accumulation of round-shaped cells with mitotic arrest, indicating the defect in mitotic spindle formation (**Supplementary Fig. 1E, F**)^42^.

**Figure 2.**
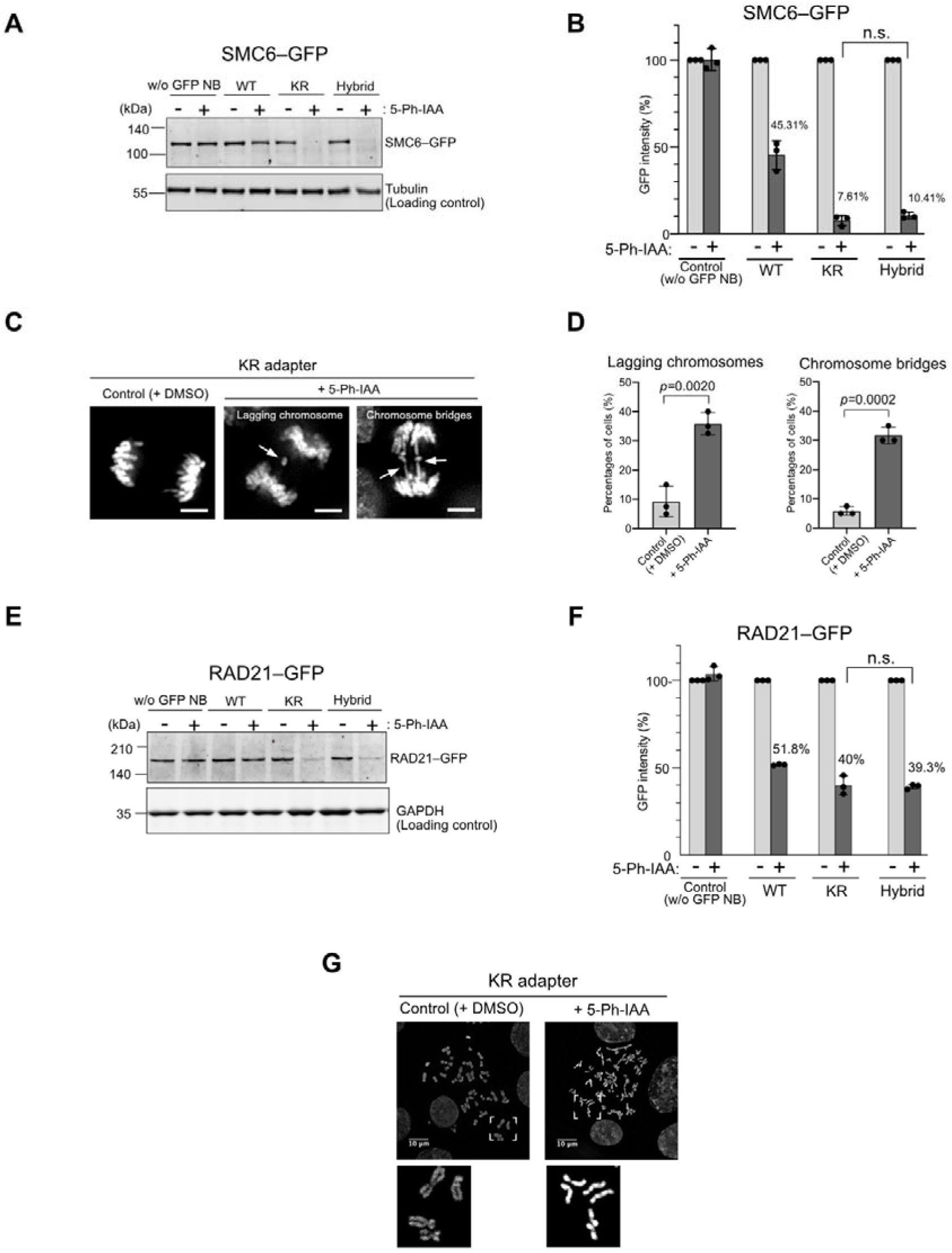
sc-AID2 with an anti-GFP nanobody adaptor enables degrading GFP-fused proteins. **(A)** An all-in-one vector shown in Fig. 1C was introduced to HCT116 cells expressing SMC6–GFP. They were treated with 1μM 5-Ph-IAA for 4 h. SMC6–GFP was detected using anti-SMC6 antibody. Tubulin was used as a loading control. **(B)** GFP intensity was analysed by flow cytometry, taking the mock-treated cells as 100%. The error bar shows the mean ± SD of three experimental replicates (n = 3). **(C)** SMC6–GFP cells expressing KR adaptor in anaphase were analysed after treatment with DMSO or 5-Ph-IAA for 24 h. Arrows indicate lagging chromosomes or chromosome bridges. Scale bar: 5 μm. **(D)** Quantification of the cell number showing lagging chromosomes or chromosome bridges. Data were presented as mean ± SD of three experimental replicates (n = 3). In total, 300 mitotic cells were analysed. **(E)** RAD21–GFP was degraded in HCT116 cells expressing the indicated adaptors in the presence of 1 μ M 5-Ph-IAA for 4 h. Cells without expressing anti-GFP nanobody were used as a negative control. RAD21–GFP was detected using anti-RAD21 antibody. GAPDH was used as a loading control. **(F)** GFP intensity was analysed by flow cytometry, taking the mock-treated cells as 100%. The error bar shows the mean ± SD of three experimental replicates (n = 3). **(G)** Spread mitotic chromosomes were visualised after treating the KR adaptor-expressing cells with DMSO or 1 μM 5-Ph-IAA for 6 h. Scale bar: 10 μm. All *p*-values in this figure were calculated using a two-tailed unpaired *t* test.

We were curious about the differences in depletion levels of the GFP-fused proteins shown above. To study the difference at the single-cell level, we employed flow cytometry to investigate the expression levels of SMC6– or RAD21–GFP with the KR adaptor in each cell (**Supplementary Fig. 2A**). We found that the KR adaptor expression levels were similarly variable over 100 folds in both SMC6– and RAD21– GFP cells, but the expression level of RAD21–GFP was 2.7 folds higher than that of SMC6–GFP. In SMC6–GFP cells treated with 5-Ph-IAA, SMC6–GFP was depleted in cells with various KR adaptor levels (**Supplementary Fig. 2A, SMC6–GFP**). In contrast, RAD21–GFP was depleted only in cells expressing low levels of the KR adaptor (**Supplementary Fig. 2A, RAD21–GFP**). Considering that the ratio between the adaptor and the target protein is likely crucial for degradation, this result was somewhat unexpected but suggests that high expression levels of the KR adaptor do not achieve efficient depletion, and proteins expressed at high levels are more challenging to degrade. If this was the case, we expected that the RAD21–GFP cells expressing lower levels of the KR adaptor would degrade RAD21–GFP more efficiently. To test this idea, we isolated three clones of the RAD21–GFP cells expressing a different level of the KR adaptor and tested the depletion after 5-Ph-IAA treatment (**Supplementary Fig. 2B**). In clones A and B, which expressed lower levels of the KR adaptor, RAD21–GFP depletion was more efficient, supporting our hypothesis (**Supplementary Fig. 2C**).

Next, we compared the depletion kinetics of scAb-AID2 and AID2 by focusing on SMC6 and RAD21 fused with GFP (scAb-AID2) or mAID–GFP (AID2) (**Fig. 3**)^17^. Initially, we treated the AID2 cells for SMC6 and RAD21 with varying concentrations of 5-Ph-IAA and found that DC_50_ was 0.64 and 0.93 nM, respectively (**Fig. 3A, C, AID2, dotted line**). These results are consistent with our previous report and indicate that AID2 induced robust degradation of SMC6 and RAD21^17^. We similarly treated the SMC6– and RAD21–GFP cells expressing the KR adaptor and found that DC50 was 0.64 and 391 nM, respectively (**Fig. 3A, C, scAb-AID2, solid line**).

**Figure 3.**
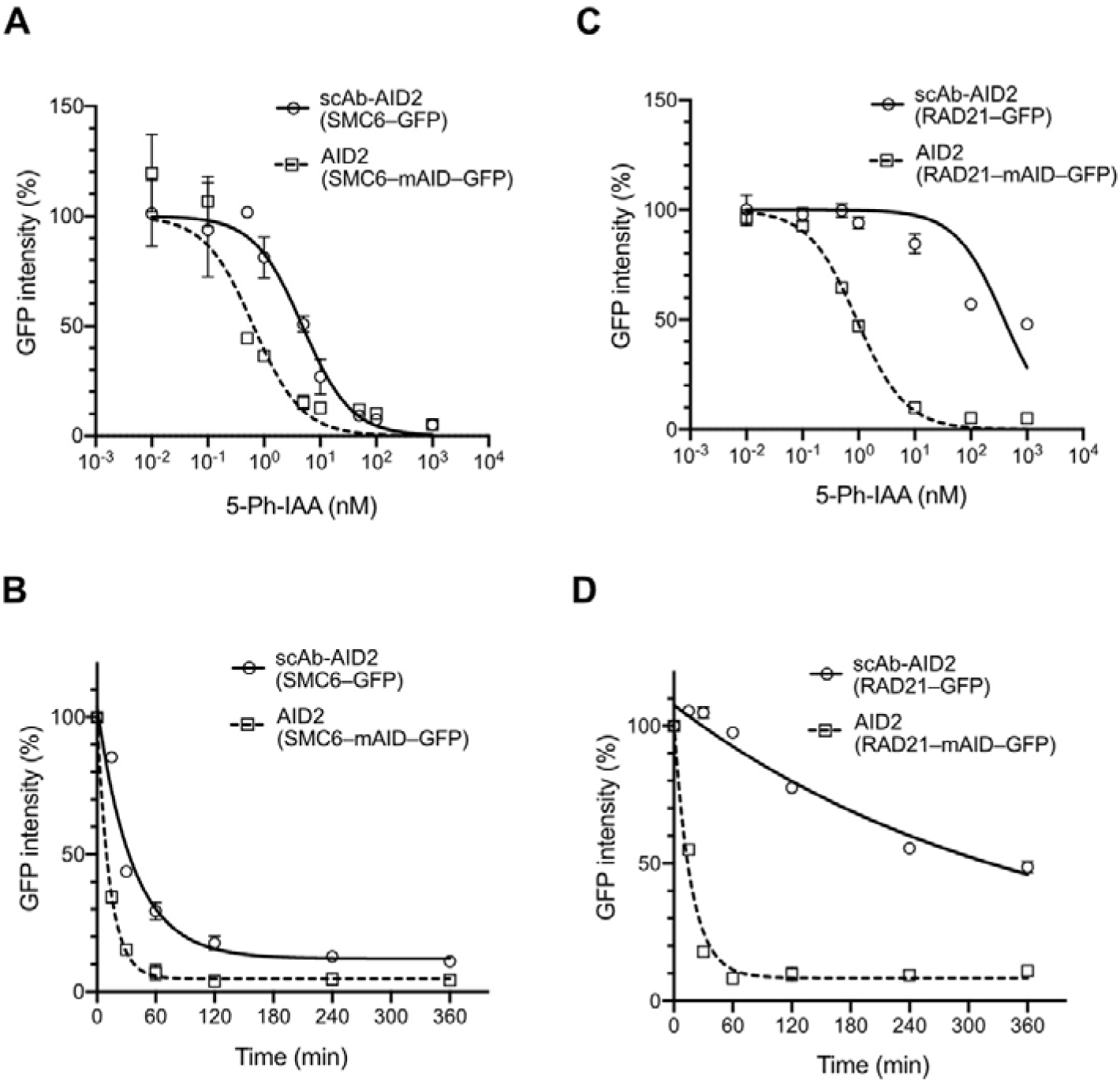
Comparison of the AID2 and scAb-AID2 systems. **(A)** Dose-response curve of SMC6–GFP depletion. The AID2 and sc-AID2 cells were treated with a range of concentrations of 5-Ph-IAA for 4 h. GFP intensity was measured using a flow cytometer, taking the mock-treated cells as 100%. The error bar shows the mean ± SD of three experimental replicates (n = 3). Data were fitted using a non-linear regression model. **(B)** Depletion kinetics of the SMC6–GFP. The AID2 and scAb-AID2 cells were treated with 1 μM 5-Ph-IAA. The GFP intensity was measured at the indicated time points using a flow cytometer, taking the mock-treated cells as 100%. The error bar shows the mean ± SD of three experimental replicates (n = 3). The data were fitted with one-phase decay. **(C)** Dose-response curve of RAD21–GFP depletion. The AID2 and sc-AID2 cells were treated with a range of concentrations of 5-Ph-IAA for 4 h. GFP intensity was measured using a flow cytometer, taking the mock-treated cells as 100%. The error bar shows the mean ± SD of three experimental replicates (n = 3). Data were fitted using a non-linear regression model. **(D)** Depletion kinetics of RAD21–GFP. The AID2 and sc-AID2 cells were treated with 1 μM 5-Ph-IAA. The GFP intensity was measured at the indicated time points using a flow cytometer, taking the mock-treated cells as 100%. The error bar shows the mean ± SD of three experimental replicates (n = 3). The data were fitted with one-phase decay.

These results indicate that scAb-AID2 works less efficiently than AID2, particularly in degrading RAD21. We then investigated depletion kinetics by treating these cells with 1 µM 5-Ph-IAA. In the case of AID2, T1/2 was 9.1 and 12.1 min for SMC6 and RAD21, respectively (**Fig. 3B, D, AID2, dotted line**). In the case of depleting SMC6– and RAD21-GFP using scAb-AID2, T1/2 was 25.6 and 243.6 min, respectively (**Fig. 3B, D, scAb-AID2, solid line**). Taking all results and the action of target recognition by the adaptor, we concluded that target depletion by scAb-AID2 is less efficient than by AID2, and depletion efficiency can be variable depending on the target protein.

### scAb-AID2 mediated protein knockdown in *C. elegans*

The original AID system has successfully been applied to *C. elegans*^43^, and we and others have recently reported that AID2 using AtTIR1(F79G) could achieve a sharper target degradation^20,21^. We have also demonstrated that membrane-permeable 5-Ph-IAA acetoxymethyl ester (5-Ph-IAA-AM) exhibits more efficacy in the embryos because it is more eggshell permeable than 5-Ph-IAA^20^. Taking these technical advances of inducible degradation in *C. elegans*, we wondered whether scAb-AID2 could be applied to degrade GFP-fused proteins in *C. elegans*. To test this idea, we used a worm line expressing the β-catenin homolog, WRM-1, fused with GFP (**Supplementary Fig. 3A, *wrm-1::GFP***). Subsequently, we introduced a transgene expressing AtTIR1(F74G) and a hybrid adaptor containing two copies of the anti-GFP nanobody under the control of the *mex-5* promoter, which is activated in germline and early embryos (**Supplementary Fig. 3A, *Pmex-5::AtTIR1(F79G)::F2A::GFP hybrid adaptor***). We grew the nematode on an agar plate containing 5-Ph-IAA-AM, transferred the embryo for observation and quantified WRM-1::GFP in the nucleus (**Fig. 4A, dotted circles**). We found that the expression level of nuclear WRM-1::GFP became 36.6% in the presence of 5-Ph-IAA-AM compared to control (**Fig. 4B**). The laid eggs did not hatch and eventually died in the presence of 5-Ph-IAA, consistent with WRM-1’s essential role in endoderm formation (**Fig. 4C, D**)^44^. This lethal phenotype was observed only in the nematodes expressing the adaptor in the presence of 5-Ph-IAA, indicating that neither the adaptor expression nor 5-Ph-IAA addition caused lethality (**Supplementary Fig. 3B, C**). These results demonstrated that scAb-AID2 works in *C. elegans*.

**Figure 4.**
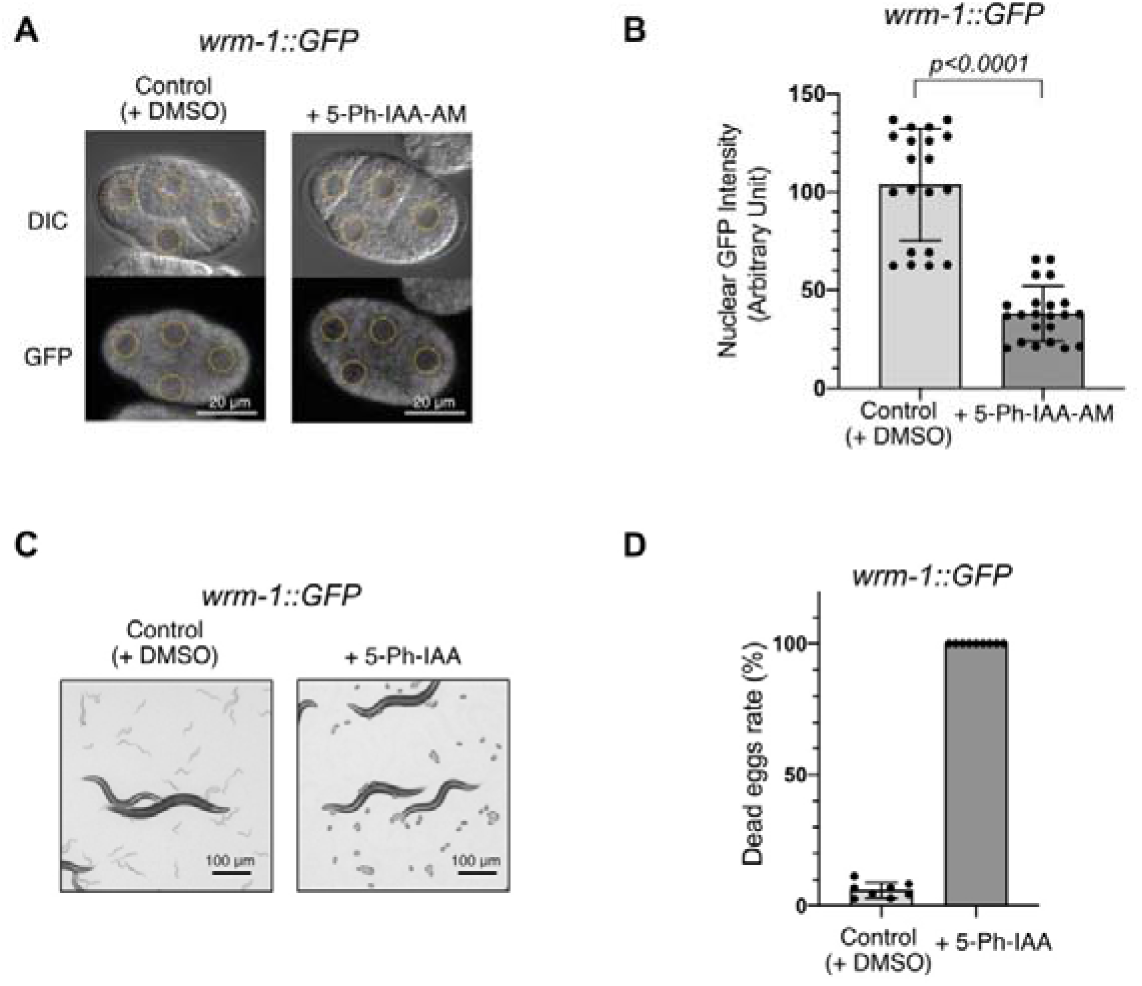
The sc-AID2 system enables degrading GFP-fused WRM-1 in *C. elegans*. **(A)** Representative confocal images of 4-cell-stage embryos expressing WRM-1::GFP, which were grown in the presence of DMSO or 5-Ph-IAA-AM. Nuclei are indicated with dotted circles. Scale bar: 20 μm. **(B)** Graph showing GFP intensity within the nuclei in the 4-cell-stage embryos treated with DMSO or 5-Ph-IAA-AM. The signal was normalized by subtracting the background signal obtained from embryos of a strain lacking WRM1-GFP expression. The error bar shows mean ± SD (n = 20 and 22 for DMSO and 5-Ph-IAA, respectively). *p*-value in this figure was calculated using a two-tailed unpaired *t* test. **(C)** Images of the sc-AID2 strain. Hatched larvae and unhatched eggs were visible in DMSO and 5-Ph-IAA plates, respectively. **(D)** Percentage of dead eggs, taking the total hatched larvae and dead eggs as 100%. A total of over 1,200 hatched larvae and dead eggs were quantified from 8 and 9 plates treated with DMSO and 5-Ph-IAA, respectively. The error bar shows the mean ± SD.

### Protein knockdown of untagged endogenous proteins

Next, we were interested in degrading endogenous proteins without fusing any tag. For this purpose, we utilized a nanobody against p53 (**Fig. 5A**)^45^. Initially, we introduced a co-expression plasmid encoding a KR or Hybrid adaptor against p53, selected a stable cell population, and subjected them to the treatment with 1 µM 5-Ph-IAA. The expression levels of p53 became about 42.3% or 47.3% in the cells expressing the KR or Hybrid adaptor, respectively, within 6 h (**Fig. 5B, C**). The p53 protein is crucial in maintaining cell viability when replication stress accumulates, and p53 knockout (p53 KO) cells show high sensitivity to aphidicolin, which induces fork stalling and replication stress (**Fig. 5D, E, compare WT and p53 KO**)^46^. We tested the viability of cells expressing the p53 KR adaptor in the presence of aphidicolin. The cells without 5-Ph-IAA showed similar aphidicolin sensitivity to WT, indicating that the expression of the adaptor did not interfere with the p53 function (**Fig. 5D, E**). In contrast, the cells treated with 5-Ph-IAA showed higher sensitivity to aphidicolin, though their sensitivity was not as high as the p53 KO cells. This is consistent with the level of p53 depletion by scAb-AID2 (**Fig. 5B, C, KR**).

**Figure 5.**
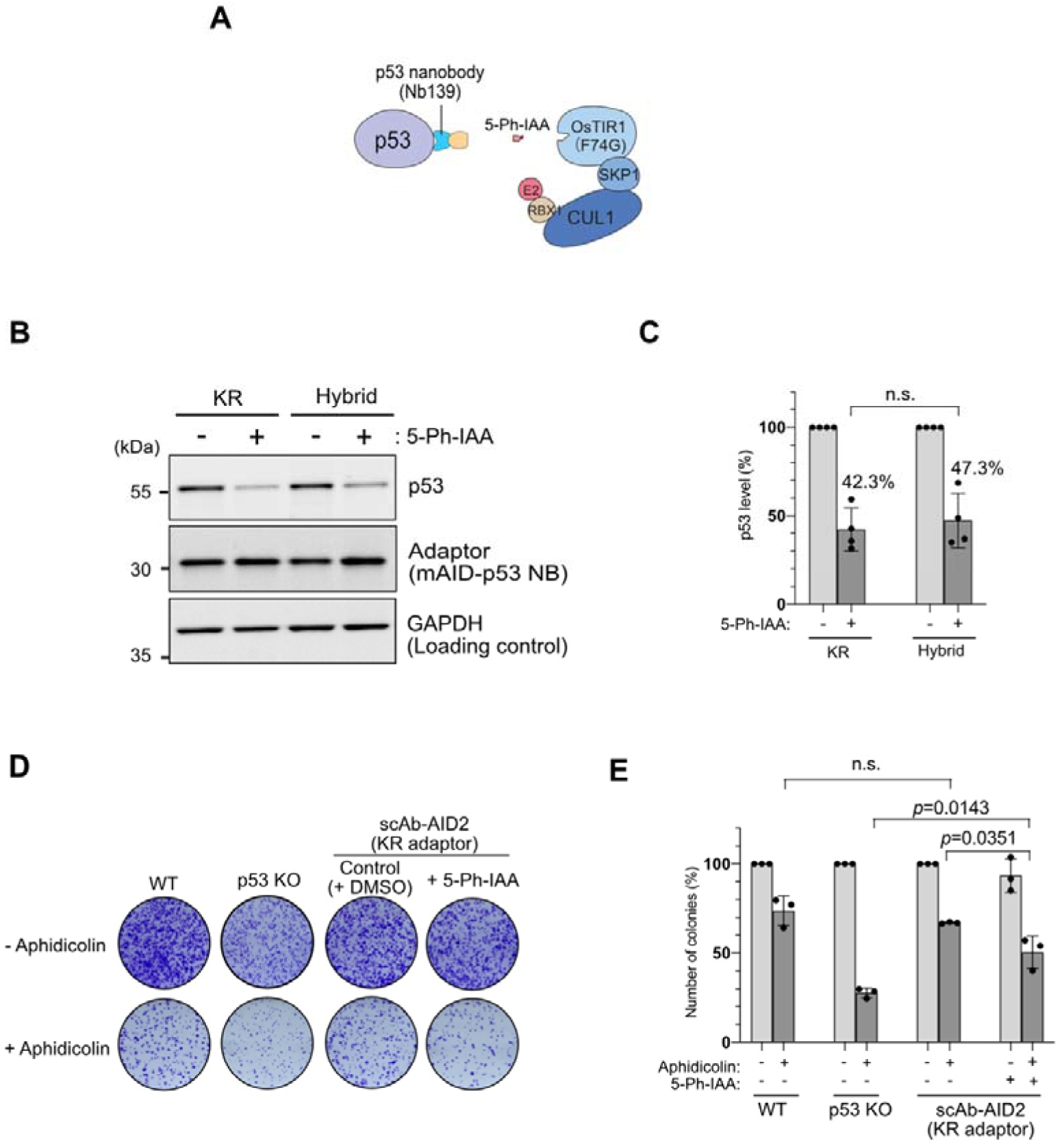
Depletion of the endogenous p53 protein using sc-AID2. **(A)** Schematic illustration of endogenous p53 protein targeted by sc-AID2. **(B)** HCT116 cells expressing KR or Hybrid adaptor were treated with or without 1 μM 5-Ph-IAA for 6h. Endogenous p53 protein, adaptor and OsTIR1(F74G) proteins were detected by Western blotting using anti-p53, -HA and -OsTIR1 antibodies, respectively. GAPDH was used as a loading control. **(C)** The graph shows the quantified p53 levels from the blot data shown in panel B. The error bar shows mean ± SD from experimental replicates (n = 3). **(D)** Colony formation of the HCT116 cells expressing the KR adaptor composed of mAID–p53 nanobody. Cells were treated with or without 1 μM aphidicolin in the presence of DMSO or 1μM 5-Ph-IAA for 24 h. Subsequently, the treated cells were seeded in a six-well plate. Colonies were stained with crystal violet after 7 days. **(E)** The graph shows the viability in the indicated conditions, taking the untreated cells as 100%. The error bar shows the mean ± SD from experimental replicates (n = 3). *p*-values were calculated using a two-tailed unpaired t-test.

Following the above strategy, we utilized a synthetic monobody binder against H- and K-RAS (**Fig. 6A**)^47^. We introduced an all-in-one vector encoding an adaptor and OsTIR1(F74G) to HCT116 and selected a stable cell population without clone isolation. We treated the cells with 5-Ph-IAA and confirmed that both H- and K-RAS were depleted down to about 40% for H-RAS and 65% for K-RAS (**Fig. 6B** and **Supplementary Fig. 4A, B**). H- and K-RAS control the activity of the downstream MAP kinase ERK1/2 and promote cell proliferation and migration^48^. We investigated ERK1/2 phosphorylation and found that it was significantly reduced in cells depleted of H- and K-RAS by scAb-AID2 (**Fig. 6C**). These results support that the RAS function was suppressed in cells treated with 5-Ph-IAA. We investigated whether the H/K-RAS knockdown would suppress tumour formation in a xenograft assay with mice. We transplanted HCT116 WT cells or the H/K-RAS KR adaptor-expressing cells under mouse skin. One week later, we began injecting 10 mg/kg of 5-Ph-IAA, as illustrated in **Supplementary Fig. 4C**^17^. In the control experiments with HCT116WT cells, no tumour suppression was observed with or without 5-Ph-IAA, showing that 5-Ph-IAA did not affect the control tumour growth (**Supplementary Fig. 4D**). In contrast, we observed significant tumour suppression when 5-Ph-IAA was administered to mice transplanted cells expressing the anti-RAS KR adaptor (**Fig. 6D, E**). Given that the RAS-ERK pathway is attenuated in the H/K-RAS KR adaptor-expressing cells (**Fig. 6C**), this is consistent with the previous report showing ERK inhibitor treatment suppressed xenograft tumour formation^49^.

**Figure 6.**
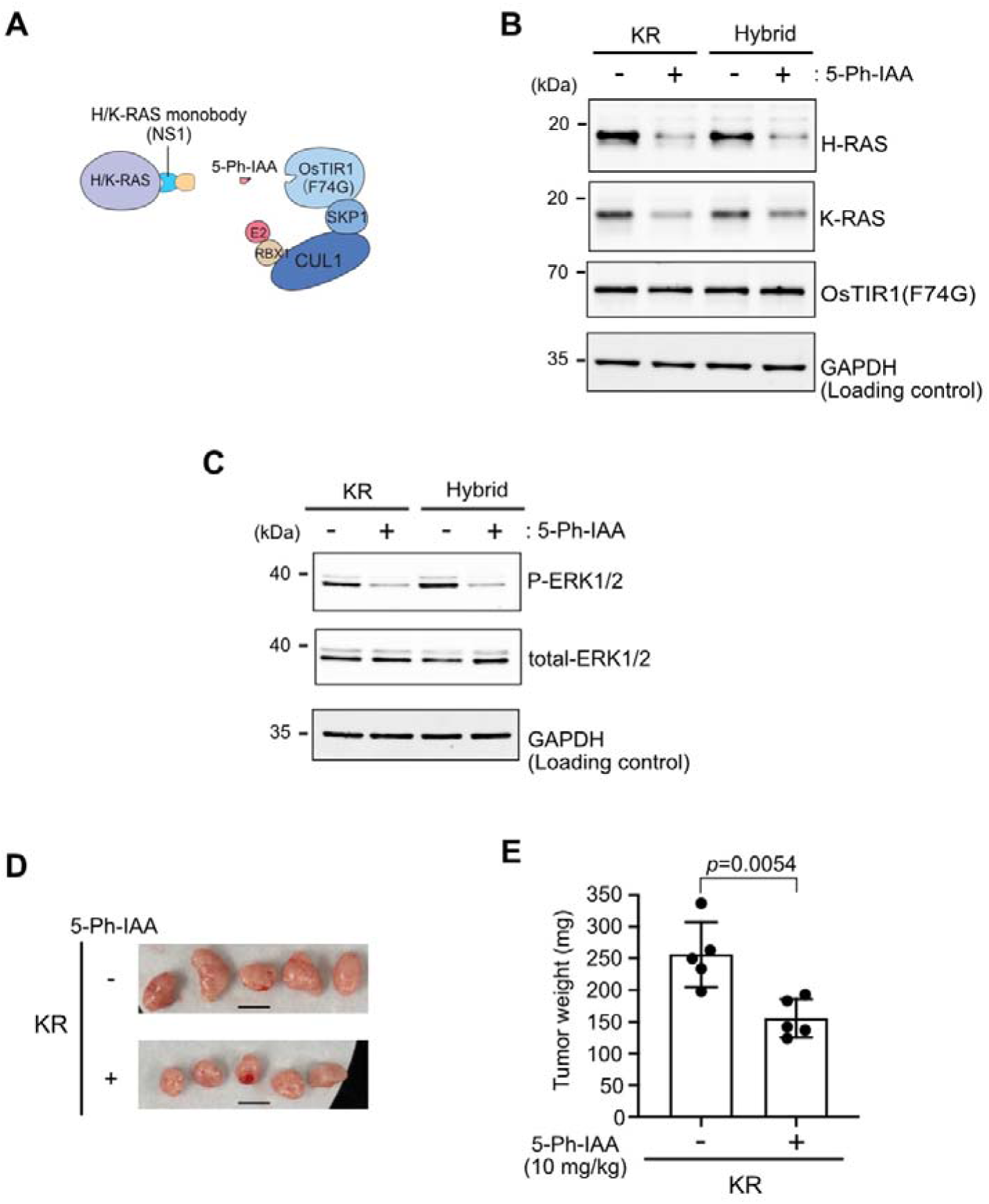
Depletion of the endogenous H/K-RAS proteins using sc-AID2. **(A)** Schematic illustration of the endogenous H/K-RAS proteins targeted by sc-AID2. **(B)** HCT116 cells expressing a KR or Hybrid adaptor were treated with or without 1 μM 5-Ph-IAA for 6 h. Endogenous H-RAS, K-RAS and OsTIR1(F74G) proteins were detected by Western blotting using anti-H-RAS, -K-RAS and -OsTIR1 antibodies, respectively. GAPDH was used as a loading control. **(C)** HCT116 cells expressing the KR or Hybrid adaptor were treated with or without 1 μM 5-Ph-IAA for 24 h. Phosphorylated ERK1/2 and total ERK1/2 were detected by Western blotting using phospho-specific ERK1/2 or pan ERK1/2 antibodies, respectively. GAPDH was used as a loading control. **(D)** Xenograft tumours of HCT116 cells expressing the KR adaptor on day 15. The scale bar shows 1 cm. **(E)** Weight of the xenograft tumours on day 15. Data are presented as mean values ± SD (n = 5 animals). The *p*-value was calculated using a two-tailed unpaired t-test.

Overall, **Fig. 5** and **6** demonstrated that both p53 nanobody and RAS monobody recognised the endogenous target protein and induced degradation by scAb-AID2. The deletion of p53 resulted in profound sensitivity to aphidicolin in cell cultures (**Fig. 5D, E**). The depletion of H/K-RAS resulted in the attenuation of the RAS-ERK pathway (**Fig. 6C**) and inhibited the growth of xenograft tumours (**Fig. 6D, E**). These results indicate that endogenous proteins can be conditionally degraded and lead to phenotypic defects using scAb-AID2.

## Discussion

It has been shown that fusing a single-chain antibody, such as an anti-GFP nanobody, to an E3 ubiquitin ligase induces degradation of GFP-fused proteins or other binding proteins (e.g. deGradFP, AdPROM, bioPROTAC)^31,33,35,50,51^. While these methodologies are useful, the target protein continues to be degraded as long as the fusion ligase is expressed, and they do not show a sharp conditional control. In contrast, there are only a few methodologies utilizing a single-chain antibody showing conditional control^36,37^. The scAb-AID2 methodology shows great temporal control (**Fig. 3C**), and target proteins are depleted within a few hours in many cases. Additionally, it can be reversibly controlled (**Supplementary Fig. 1A, B**) and works not only in mammalian cells but also in nematodes (**Fig. 4**). Taking into account that AID2 can be used in many eukaryotic species^17,20,52^, scAb-AID2 has great potential to aid basic research.

We have shown that GFP-fused proteins can be induced to degrade by utilizing a GFP-nanobody adaptor in mammalian cells and *C. elegans*, as previously demonstrated in mammalian cells with AID and in yeast with AID2/ssAID (**Figs. 2** and **4**)^36,38^. As far as we know, this is the first demonstration that an antibody-mediated conditional degradation technology worked in *C. elgans*. Since the Cas9- mediated knock-in became available, there have been many GFP cell lines and *C. elegans* lines have been reported^53–55^. The scAb-AID2 technology described in this paper will be useful to knock down GFP-fused proteins in the existing cell and *C. elegans* lines. We also suggest that scAb-AID2 can be applied to other GFP- expressing organisms, such as yeasts, *Drosophila* and zebrafish^56–61^. By utilising the established GFP lines, researchers can efficiently implement a degradation strategy by introducing an all-in-one vector expressing OsTIR1(F74G) and a GFP nanobody adaptor.

We noted that our GFP-nanobody adaptor does not show localization preference and changes its location depending on the target protein fused with GFP (**Supplementary Fig. 5**). Therefore, the GFP-nanobody adaptor binds constitutively to the target in either the nucleus or cytoplasm. Users should note that the adaptor binding might interfere with the function of bound proteins, even when degradation was not induced.

Not only to degrade GFP-fused proteins, we have shown that endogenous proteins, p53 and H/K-RAS, can be induced for degradation in human cells and succeed in showing their phenotypic changes upon degradation (**Figs. 5** and **6**). In principle, it is now possible to generate a single-chain antibody or a synthetic binding peptide to any proteins of interest^29,62,63^. Therefore, scAb-AID2 opens a new avenue to degrade untagged endogenous proteins in various organisms.

Although scAb-AID2 works in mammalian cells and *C. elegans*, we found that its depletion efficiency is lower and target-dependent compared to AID2 (**Fig. 3**). While AID2 has shown greater depletion efficacy for many proteins, its application in a therapeutic context is constrained due to the challenges involved in tagging target proteins. One of the major benefits of scAb-AID2 is that it does not require genetic manipulation to introduce degron tags directly onto target proteins. As the field of single-chain antibodies grows rapidly^29,62,63^, leveraging non-invasive single-chain antibodies allows researchers to circumvent the challenges associated with fusing a degron tag to a target protein. Even though chemical heterobifunctional degraders (e.g. PROTAC, SNIPER) can degrade untagged endogenous proteins^11,64^, developing functional degraders involves a time- and cost-consuming laborious process. Moreover, chemical degraders are unavailable for proteins to which a specific chemical binder has not been identified. Taking these facts into account, scAb-AID2 can potentially be an alternative to chemical degraders for conditionally degrading target proteins through the transfection of an all-in-one vector and the use of the inducer, 5-Ph-IAA. We provided a framework of how ligand-inducible protein interactions can be made possible by single-chain antibodies without requiring the target to be genetically modified, broadening the potential uses beyond basic research.

## Materials and Methods

### Plasmids

All plasmids used in this study are listed in **Supplementary Table 1**. They are available from Addgene.

### Cell Culture

HCT116 cells were cultured in McCoy’s 5A (Gibco, #16600-108), supplemented with 10% FBS, 2 mM L-glutamine, 100 U/ml penicillin and 100 μg/ml streptomycin at 37 °C in 5% CO_2_. For generating stable cell lines expressing an adaptor and OsTIR1(F74G), an all-in-one plasmid shown in **Fig. 1B** and pCS-TP encoding a Tol2 transposase was co-transfected using ViaFect (Promega, #E498A) and Opti-MEM (ThermoFisher Scientific, #31985062) in a 12-well plate following the manufacturer’s instruction^65,66^. Stable cells were selected in the presence of 100 μg/mL Hygromycin B Gold (InvivoGen, #ant-hg-1). Cells were treated with DMSO or 5-Ph-IAA for the indicated time. 5-Ph-IAA was synthesized as previously described^17^. All cell lines used in this study are summarized in **Supplementary Table 2**.

### Protein Detection

Cells were seeded at 2×10^5^ cells/well in a 6-well plate and grown for 2 days. Cells were lysed in RIPA buffer (25 mM Tris-HCl pH7.6, 150 mM NaCl, 1% NP40, 1% sodium deoxycholate, 0.1% SDS) containing complete protease inhibitor cocktail (Roche, #1187580001) for 30 min on ice. Cells were then centrifuged for 20 min at 4 °C. The supernatant was mixed with 2 × SDS sample buffer (Cosmo Bio, #423420) before incubation at 95 °C for 5 min. An equal amount of protein samples was loaded on a 7.5% or 10% TGX Stain-Free gel (BioRad) or 12% SDS polyacrylamide gel. Separated proteins were transferred to a nitrocellulose membrane (Cytiva, #10600003) using a semi-dry Trans-blot Turbo system (BioRad) or a wet/tank blotting system (BioRad).

The membrane was processed with 1% skim milk in TBS-T for 30 min or 2% ECL Prime blocking agent (Cytiva, RPN418V) for 1 h at room temperature. For phosphorylated ERK and ERK detection, the membrane was processed with 1% BSA/TBST for 30 min at room temperature. The membrane was incubated with a primary antibody at 4 °C overnight and subsequently incubated with a secondary antibody at room temperature for 1 h. For protein detection antibodies were diluted with TBST containing 5% skim milk or ECL blocking buffer or 1% BSA/TBST. Proteins were detected by the ChemiDoc Touch MP imaging system (Bio-Rad), and the detected signals were quantified using Image Lab software (Bio-Rad). The data shown are representative of at least three biological replicates.

### Antibodies

All antibodies used in this study are listed in **Supplementary Table 3**.

### Microscopy

To observe mitotic chromosomes after SMC6–GFP depletion (**Fig. 2C, D**), cells growing on coverslips were incubated with 20 µM lovastatin for 20 h to arrest them in the G1 phase. One µM 5-Ph-IAA was added at 16 h after the addition of lovastatin.

The G1-arrested cells were released by washing three times with prewarmed fresh media and then were allowed to enter the S phase in the presence of 1 µM 5-Ph-IAA and 2 mM mevalonate. The released cells were re-arrested in the G2 phase by adding 7 µM RO-3306 at 17 h after release and incubated for an additional 3 h. To induce anaphase, the G2-arrested cells were released for 40 min before fixing the cells. Cells on coverslips were fixed with 4% paraformaldehyde and then permeabilized with PBS containing 0.1% Triton X. Cells on the coverslip were mounted with VECTASHIELD anti-fade mounting medium (Vector Laboratories, #H-1000-10) containing DAPI. Images were observed under a DeltaVision deconvolution microscope (GE Healthcare).

To observe mitotic chromosome cohesion after RAD21–GFP depletion (**Fig. 2G**), cells were cultured to 70% confluency in a 60-mm dish. KaryoMAX Colcemid Solution in PBS (Gibco, #15212012) was added to a final concentration of 0.02 μg/ml together with DMSO (control) or 1 μM 5-PhI-AA. Treated cells were incubated at 37 °C in 5% CO_2_ for 6 h before trypsinization. Removed cells were treated with 75 mM KCl before fixation in MeOH/acetic acid (3:1). Fixed cells were adjusted to approximately 10^7^ cells/ml. Ten μl of the cell suspension was applied onto a glass slide and dried at room temperature. 10 μl of DAPI-containing Vectashield Mounting Medium (Vector Laboratories, #H-1200) was added before sealing with a coverslip. Images were observed under a DeltaVision deconvolution microscope (GE Healthcare).

For fluorescent microscopic images (**Supplementary Fig. 5**), cells on coverslip were fixed with 4% paraformaldehyde in PBS for 15 min at room temperature. After washing with PBS-T (0.05% Tween-20 in PBS), the fixed cells were permeabilized with 0.5% TritonX-100 in PBS for 5 min at room temperature. Following permeabilization, cells were washed and blocked with 5 % BSA in PBS for 20 min at room temperature. After blocking, cells were washed with PBS-T and incubated with mouse monoclonal anti-HA antibodies diluted in 1% BSA in PBS for 1.5 h at room temperature. After washing with PBS-T, cells were incubated with DAPI and Alexa 594-conjugated goat anti-mouse IgG secondary antibody diluted in 1% BSA in PBS. The coverslips with cells were washed and mounted onto slide glasses with ProLong Glass antifade mount (Thermo Fisher Scientific). DAPI, GFP (mClover), and HA-tagged adaptor signals were visualized under a Nikon Ti2 microscope through CFI APO TIRF 60x/1.49 objective lens equipped with a Hamamatsu ORCA-FusionBT C15440 camera.

### Flow Cytometric Analyses

Cells were seeded at 1×10^5^ cells/well in a 6-well plate and grown for 2 days until 80– 90% confluency. For detecting GFP signals after ligand treatment, the treated cells were trypsinized and fixed in 4% methanol-free paraformaldehyde phosphate at 4 °C for 15 min. Fixed cells were washed and resuspended in PBS containing 1% BSA. Flow cytometric analysis was performed on a BD Accuri C6 flow cytometer (BD Biosciences). Ten thousand cells were analyzed from each sample after gating for single cells.

### Cell Viability Assay

Cells expressing a p53 nanobody adaptor were treated with DMSO or 1 μM 5-Ph-IAA for 6 h prior to adding 1 μM aphidicloin. After 24 h, 3000 cells were seeded in 6-well plate and cultured for 7 days. The culture medium was exchanged with a fresh medium on day 4. Cells were fixed and stained with crystal violet solution (6% Glutaraldehyde, 0.5% crystal violet). The colonies were counted using the ImageJ software.

### C. elegans Culture and Strain Construction

The strains used in this study are listed in **Supplementary Table 4**. They were cultured by standard methods^67^. To generate the strain containing the transgene encoding *Pmex-5::AtTIR1(F79G)::F2A::GFP hybrid adaptor*, CRISPR– Cas9-mediated single-copy insertion was performed as previously described^20^.

To quantify WRM-1::GFP degradation, adult *C. elegans* grown on plates containing 10 µM 5-Ph-IAA were dissected to isolate 1-cell stage embryos. These embryos were then transferred to the M9 buffer containing 100 µM 5-Ph-IAA-AM for observation. Live imaging with confocal microscopy was performed as previously described ^20^. To quantify WRM-1::GFP, its intensity was measured within the nuclei in the 4-cell-stage using Fiji^68^. The dead egg rate was determined by counting the number of dead embryos and hatched larvae on the plates containing 10 µM 5-Ph-IAA.

### Mouse Xenograft Assay

Nude mice used for xenograft assay were Balb/c-nu female mice (7-week old) weighing 16 to 20 g and were obtained from Japan SLC, Inc. (Shizuoka, Japan). These animals were acclimated at least for one week before use. Indicated HCT116 lines (4×10^5^ cells) were resuspended in 0.1 ml of HBSS (Sigma-Aldrich, #H9269) containing 0.05 ml of Matrigel (Corning, #356234). The suspension was injected into both sides of the flanks. Seven days after transplantation, the mice were randomized and treated daily with 10 mg/kg of 5-Ph-IAA by intraperitoneal (IP) injection for an additional 7 days. Tumour volume (= L×W×W/2; L and W stand for the longest and shortest diameters, respectively) was measured on the indicated days. At the end of the experiment, the xenograft tumour was removed and weighed.

### Data Analyses and Software

FCS Express Cytometry 4 (De Novo Software) and FlowJo (BD Biosciences) were used for flow cytometric analyses. Image lab software (Bio-Rad) was used for quantifying western blot data. Fiji was used for counting colonies. GraphPad Prism 6 (Dotmatics) was used for statistical analyses and graph generation. All figures were created with Affinity Designer 2 (AFFINITY).

## Acknowledgements

We thank all members of the Kanemaki laboratory for discussion and support. We also thank Tomoko Suzuki, Kosuke Yamaguchi and Yumiko Saga for their help with mouse experiments. MI is a MEXT scholarship fellow, and YH is a JSPS Research Fellow for Young Scientists (DC2). The *C. elegans* strains, N2 and RW12263, were provided by the CGC, which is funded by NIH Office of Research Infrastructure Programs (P40 OD010440). This work was supported by JSPS KAKENHI (JP23K05836) to TN, and JSPS KAKENHI (JP21H04719 and JP23H04925) and JST CREST (JPMJCR21E6) to MTK.

## Author Contributions

This project was conceived and designed by MTK with the help of the other authors. MI and YH performed experiments with cell culture. MI and TN performed experiments with *C. elegans*. NK performed mouse xenograft experiments under the supervision of MTK and KG. KH synthesised 5-Ph-IAA and 5-Ph-IAA-AM. MI and MTK wrote the manuscript with contributions from most authors.

