## Supplementary Figures 1-5 for "A single-chain antibody-based AID2 system for conditional degradation of GFP-tagged and untagged proteins"

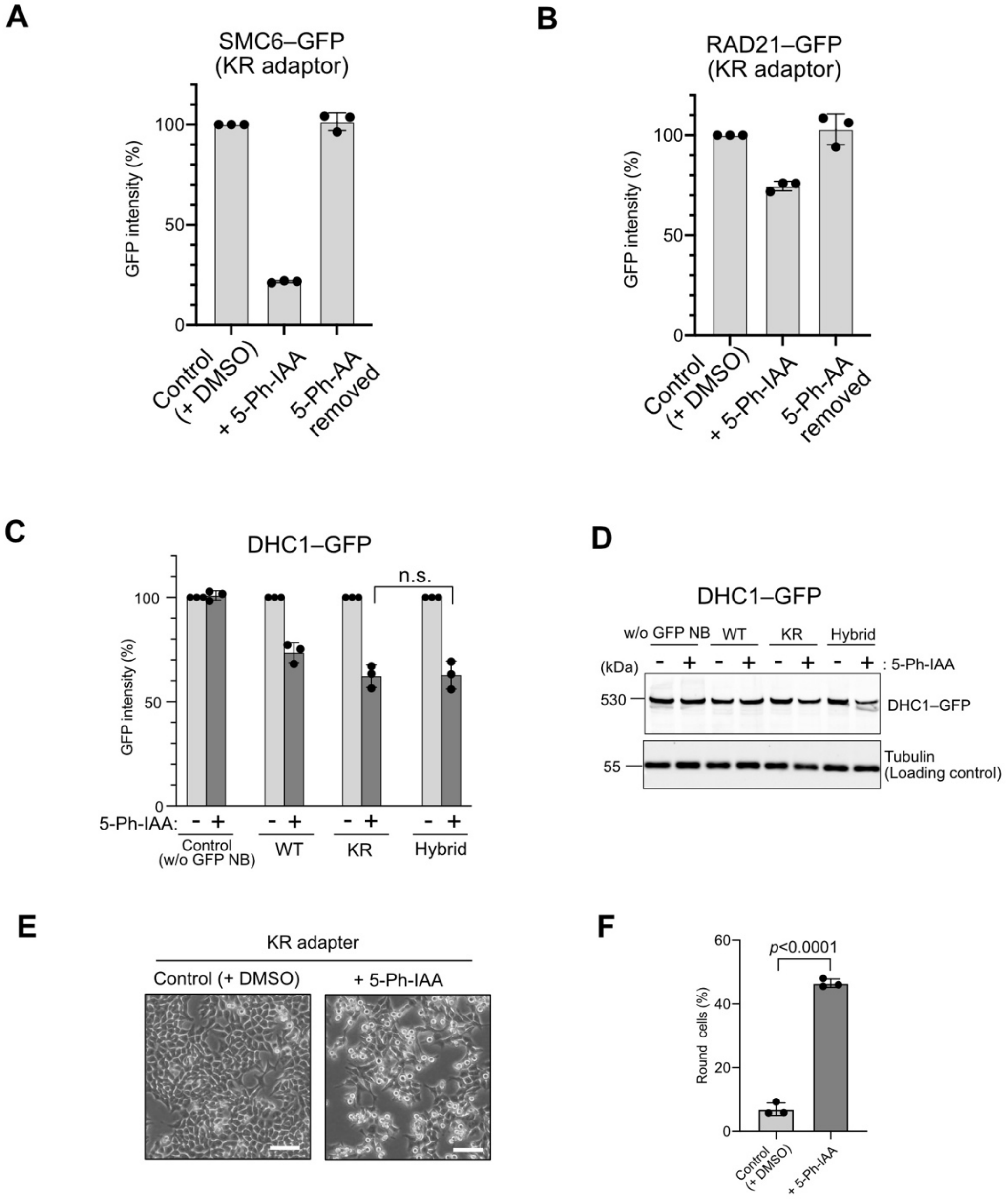


**Supplementary Figure 1**

Degradation of GFP-fused proteins in the nucleus and cytoplasm. **(A, B)** Re-expression of SMC6– and RAD21–GFP after depletion. The SMC6–GFP cells expressing the KR adaptor were treated with 50 nM 5-Ph-IAA for 4 h. Subsequently, cells were washed and cultured in a fresh medium without 5-Ph-IAA for 24h. GFP intensity was analysed using a flow cytometer, taking the mock-treated cells as 100%. The error bar shows the mean ± SD of experimental replicates (n = 3). **(C)** DHC-1–GFP was induced to degrade in HCT116 cells expressing the indicated adaptor by adding 1 μM 5-Ph-IAA for 4 h. GFP intensity was analysed using a flow cytometer, taking the mock-treated cells as 100%. The error bar shows the mean ± SD of experimental replicates (n = 3). **(D)** Detection of DHC1–GFP by Western blotting. Cells were treated as in panel C. Tubulin was used as a loading control. **(E)** Cells expressing the KR adaptor were treated with either DMSO or 1μM 5-Ph-IAA for

24 h before being visualised under a microscope. **(F)** The number of round-shaped cells was counted. In total, 300 cells were counted. The error bar shows the mean ± SD of experimental replicates (n = 3). *p*-values were calculated using a two-tailed unpaired t-test.

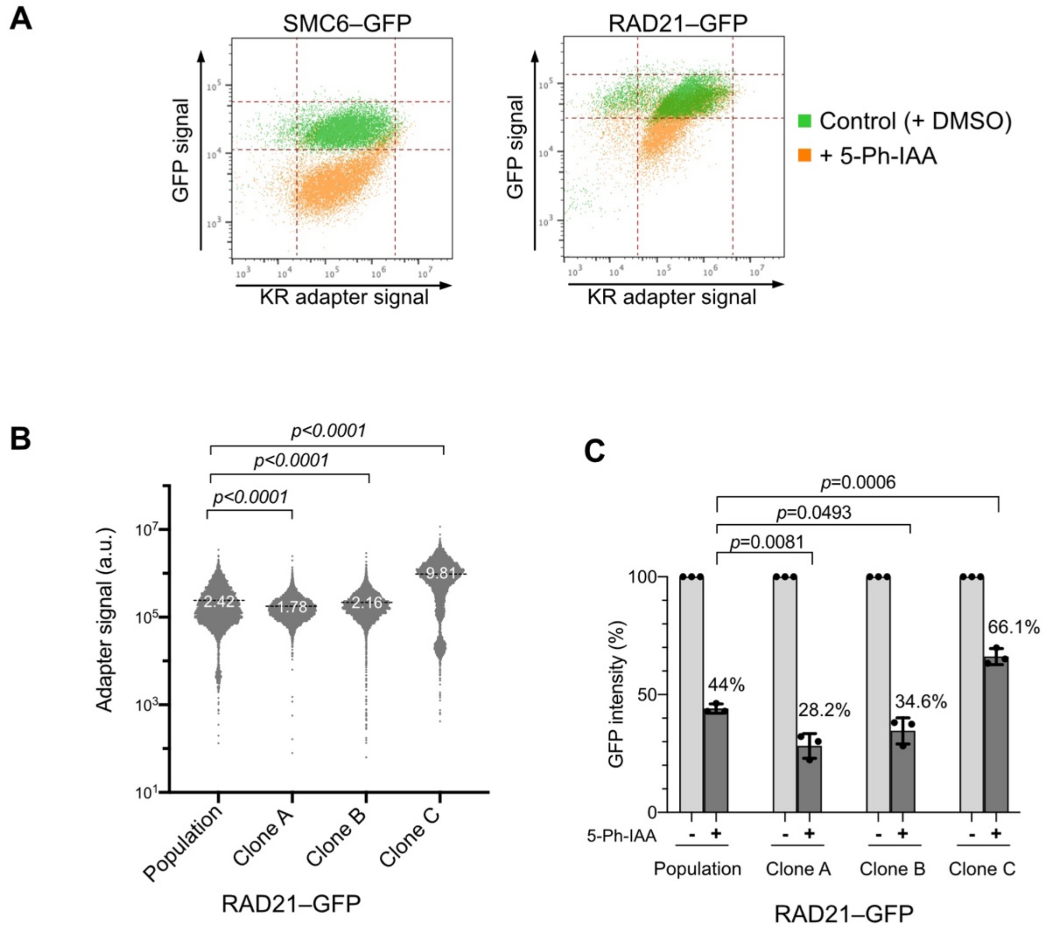


**Supplementary Figure 2**

The adapter and target protein expression levels affect the efficiency of target depletion. **(A)** Two-dimensional flow cytometry plot showing the KR adapter levels (X-axis) and the GFP-fused target protein levels (Y-axis). Green and orange dots are cells treated with DMSO and 1 μM 5-Ph-IAA for 4 h, respectively. **(B)** RAD21–GFP clones expressing the KR adapter were isolated and analysed by flow cytometry to evaluate the adaptor expression level. Plots were obtained from 5000 cells. The dotted line shows the mean intensity. **(C)** The indicated RAD21–GFP clones were induced for degradation by adding 1μM 5-Ph-IAA for 6 h. GFP intensity was analysed by flow cytometry, taking the mock-treated cells as 100%. The error bar shows the mean ± SD of experimental replicates (n = 3). *p*-values were calculated using a two-tailed unpaired t-test.

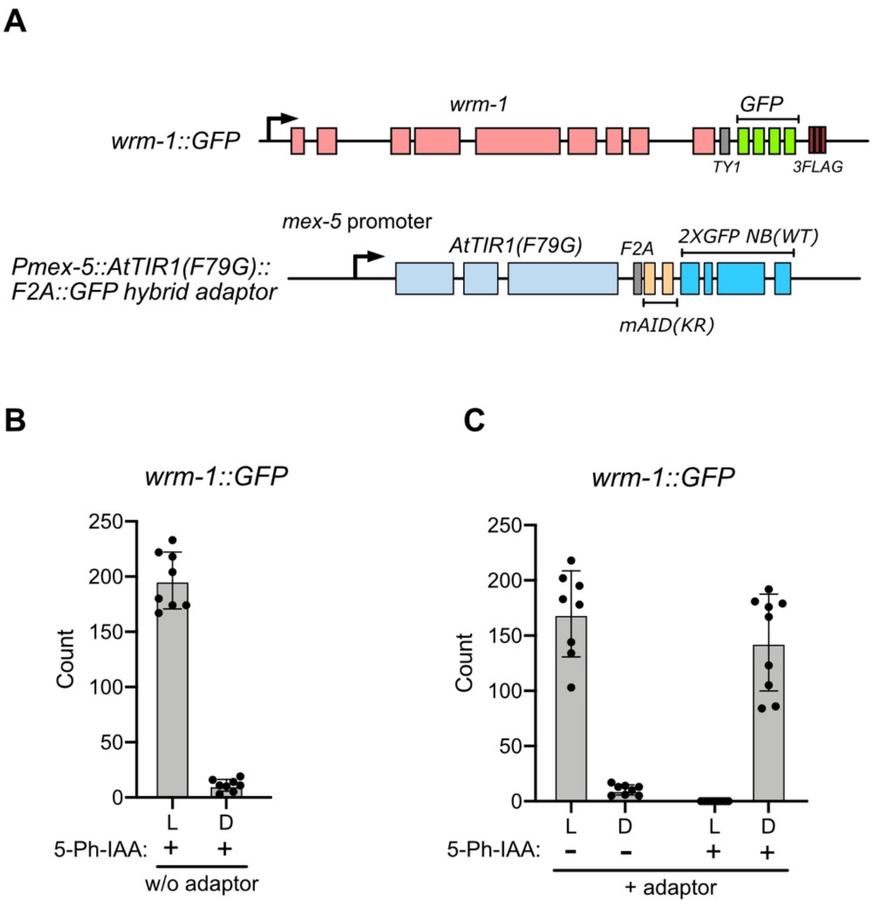


**Supplementary Figure 3**

Induced degradation of WRM-1::GFP in *C. elegans*. **(A)** Illustration of the genotype showing the nematode strain used for WRM-1::GFP degradation. **(B)** The strain with or without the adaptor was grown in the presence of 5-Ph-IAA-AM. A total of 1572 hatched larvae and dead eggs were examined. The error bar shows the mean ± SD of experimental replicates (n = 8). **(C)** The strain with the adaptor was grown in the presence or absence of 5-Ph-IAA. A total of over 1,200 hatched larvae and dead eggs were analysed. The error bar shows the mean ± SD of experimental replicates (n = 8 and 9 for DMSO and 5-Ph-IAA-AM, respectively).

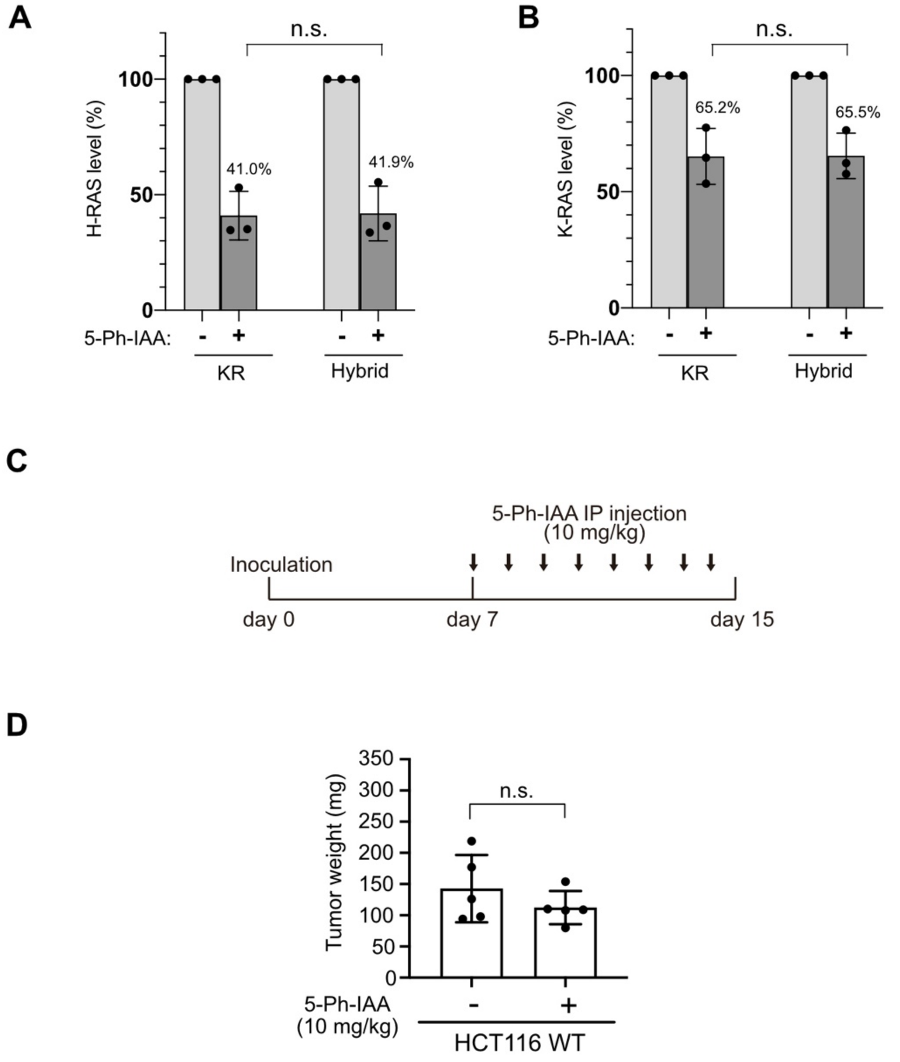


**Supplementary Figure 4**

Related to Figure 6. **(A, B)** The graphs show the expression levels of H-RAS and K-RAS proteins from the blot data in panel B. The error bar shows the mean ± SD of experimental replicates (n = 3, two-tailed t-test). **(C)** Experimental time-course diagram showing the xenograft assay. **(D)** Weight of HCT116 WT xenograft tumours on day 15. Data are presented as mean values ± SD (n = 5 animals, two-tailed t-test).

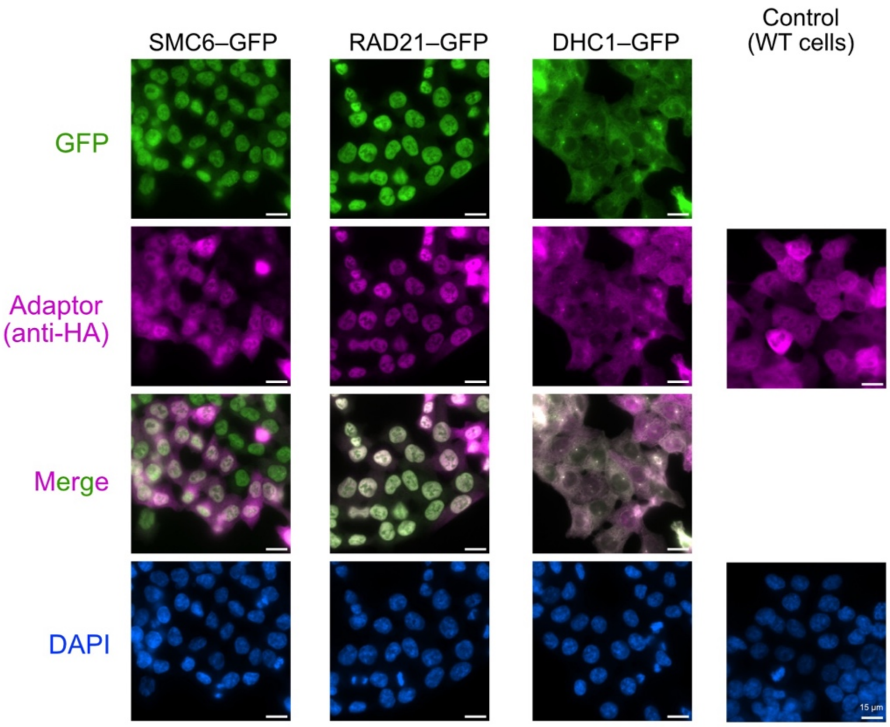


**Supplementary Figure 5**

Fluorescent microscopic images showing the cellular localisation of the KR adapter and the indicated GFP-fused proteins. Cells were stained with DAPI (blue) and anti-HA antibody (magenta). The scale bar indicates 15 μm.
